# Predation by Himalayan Wolves: Understanding conflict, culture and co-existence amongst Indo-Tibetan community and large carnivores in High Himalaya

**DOI:** 10.1101/2019.12.16.877936

**Authors:** Salvador Lyngdoh, Bilal Habib

## Abstract

The wolves in the Hindukush-Himalayan region belong to one the most basal lineages within *Canis lupus*, yet little is known about its ecology, distribution, and behavior. To understand ecological aspects of wolves in this landscape, we predict wolf distribution, diet patterns and conflict perception in Spiti, India using field and remotely sensed information. We collected scats (n = 283) of canid species namely, Wolves, and other predators over a period of 3 years (2014-17) [66]. Wolf diet constituted mostly of domestic prey (79.02 %) while wild prey constituted to 17.80% of wolf diet over the three years. Village surveys recorded only 4% of the respondents confirmed wolf presence and perceived them as a possible threat to various livestock. Over, 98% of the respondents claimed that wolves were not safe for livestock and were averse to its presence. Marginal response curves depicted the model to have positive responses to animal location, LULC, village population, village density and wolf depredation. We found perceived presence/threat distribution wolves in the area significantly differed from actual ecological presence and distribution of wolves. The Himalayan wolf is an apex flagship predator in this fragile high altitude system, whose role is intricately linked with the ecology of the region. The use of such methods can aid in understanding such aspects as well as designing effective long-term conservation strategies for the species.

## Introduction

Grey wolves are found throughout the Northern Hemisphere. In the Indian Sub-continent, these are represented by geographically isolated broadly non-overlapping (allopatric) populations [1]. One population of wolves extends from the upper Hindukush-Himalayan region of India across the two northernmost states of Himachal Pradesh and Jammu and Kashmir (Fox et al., 1992). This wolf population is well adapted to the cold environment and is known as the Himalayan wolf, *Canis lupus chanco* [2]. Himalayan wolves are found in the cold and hypoxic high altitude ecosystems of the Himalayas and the Tibetan Plateau, extending into China, Manchuria and Mongolia [3]. Their unique ancient lineage was highlighted recently via several taxonomic and evolutionary studies. However, there is limited understanding of its ecology, diet, behaviour, and habitat requirements primarily due to its cryptic nature and the harsh landscape that it occupies [4–6].

Socio - cultural aspects are crucial in planning conservation strategies. History shows that wolves originally lived almost throughout North America and Eurasia. However, the only way in which wolf population was reduced was by human control [7]. Attitudes against wolves became strongly sentimental with beginning of European settlement of the Americas. The wolf became viewed as a threat to personal safety, an object to subdue and an impediment to progress and civilization [8]. Later, depredation of livestock led to government aided ‘legal’ extirpations across many areas in the United states and Canada. In Europe, wolves have been illegally hunted. In 1966, the wolf was declared functionally extinct in the Scandinavian Peninsula. [9]. Nearly, all mortalities in Scandinavia, Italy, Germany and England were reported from poaching or vehicle strikes [7].

In, the Asian context, wolves seemed to prey largely on domestic livestock [2,10,11]. Probable reasons are, reduced availability of wild prey and changing land-use into livestock grazing grounds leading to increased conflicts and retaliations towards wolves [12]. In India, however, attitudes towards wolves have been less destructive. Unlike, many western notions, wildlife and wolves have been tolerated in India due to many religio-cultural sentiments. Yet, persecution remains as one of the biggest obstacles to wolf recovery around the world [13], including the Himalayan wolf [10,14]. In many Indian landscapes, wolves continue to survive *vis-a-vis* disturbance and other human induced factors [15]. Tolerance of communities has aided survival and depredation is seen to be part of the occupation [10,16]. With unavailability of wild prey and natural habitats wolves now subsist mainly on livestock and continue to live outside protected areas [17].

There have been several studies on wolves worldwide, however the Himalayan wolf is one of the least studied wolf in spite of its genetic distinction and ancient lineage [2,18,19]. The current study thus aims to understand aspects of Himalayan wolf ecology in context of human-wolf interaction with the use of telemetry, scatological and distribution modelling techniques [20–22]. We investigate a) dietary choices of Himalayan wolves in a one such landscape and their correlates b) we try to understand wolf conflict hotspots through the study of their movement, diet and public perception. And, finally we describe prevailing religio- cultural aspects of human-wolf interactions to better manage wolf populations in Trans-Himalayan landscapes. We believe the study will add to existing information on wolf ecology particularly from the Himalayan ranges.

## Study Area and Methods

The Spiti catchment area (Fig. 1) which is divided by the Spiti River is a cold semi-arid region of the state of Himachal Pradesh. The greater Himalayas are to the south east of the valley while on the north of it lays Ladakh and east lays Tibet. The study area comprises two protected areas of Kibber Wildlife Sanctuary (WLS) (32°5 to 32°30 N and 78°1 to 78°32E) and Pin Valley National Park (NP) in the south west corner of the Spiti Catchment area. Kibber WLS was established in the Spiti Region (Lahaul and Spiti District) of the Indian state of in 1992 for conservation of trans-Himalayan wildlife. It is a mountainous cold desert, where altitudes range between 3600 m and 6700 m above mean sea level. Temperatures range between 23°C and 3°C in winter and between 1°C and 28°C in summer. The vegetation of this area has been broadly classified as dry alpine steppe [23].

**Fig 1.**
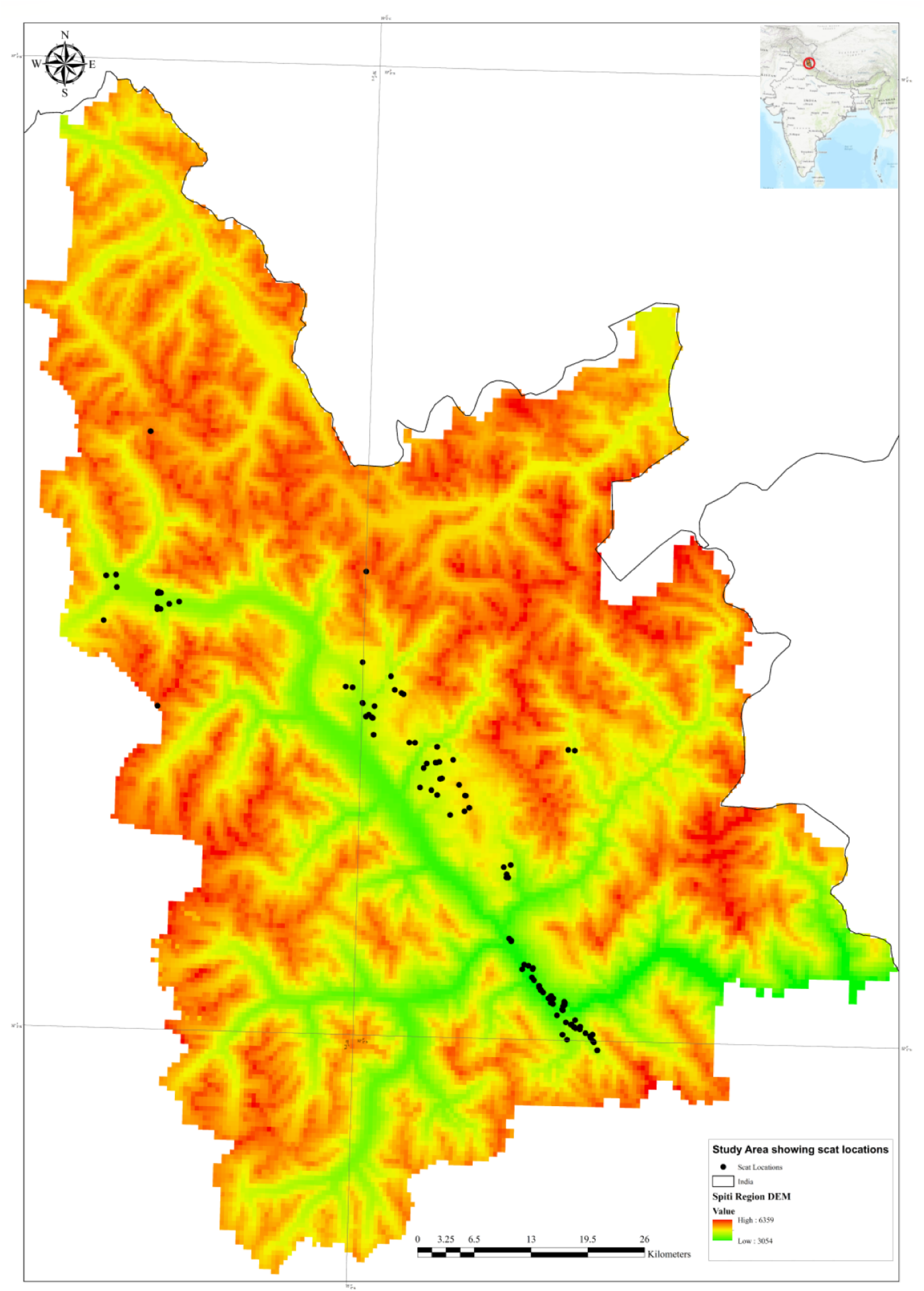
Study Area depicting Spiti region in North India with locations of scats

Large mammalian fauna of the area includes bharal (*Pseudois nayaur*), relatively few ibex (*Capra ibex*), and their predators, namely the snow leopard and the wolf. Other fauna includes the red fox, *Vulpes vulpes*, pale weasel (*Mustela altaica*), stone marten (*Martes foina*) and the Himalayan mouse hare (*Ocho tonas*). Nearly 45 species of birds were identified, including several species typical of this alpine habitat, such as chukar (*Alectoris chukar*), and Tibetan snowcock (*Tetraogallus tibetanus*).

The local agro-pastoral community, largely Buddhist, are concentrated in village clusters largely along the main Spiti River. There are approximately 60 villages through the length and breadth of the valley. Most agriculture- related activities are restricted to the short growing season (from May to September). The main crops cultivated are barley (*Hordeum vulgare*) and green peas (*Pisum sativum*). Livestock includes goats and sheep, cattle, ‘dzomo’ (a female hybrid of cattle and yak), and yaks. Donkeys are the beasts of burden, while horses, apart from being used for religious ceremonies, are mainly raised for trade. The region has traditionally practiced systems of grazing whereby nomadic as well as resident herders frequent this landscape. In this vast landscape (~ 4000, [24] the study was concentrated much towards higher elevation regions of the area (> 3000 m) from villages surrounding Pin valley NP and Kibber WLS.

## Data collection

### Village surveys

In order to determine livestock depredation and large carnivore presence we sampled the landscape by conducting questionnaire surveys (Fig.1) using a semi structured questionnaire [25] among the locals across 35 villages. The surveys were distributed systematically to represent most of the landscape. A sighting of wolf was confirmed by showing them photographs of different canids and asking details of the sighting incident and place. Any ambiguity in the confirmation was not recorded. In total 200 households were surveyed with questions mainly focussed on household characteristics such as demography, income, grazing distance, livestock holding, depredation and carnivore presence.

### Data on wolf food habits

Sign surveys were conducted throughout the landscape with more than 1000 km tracked and 300 man days of effort. Trails, river banks, hill tops, village periphery and grazing pastures were searched for signs of a large predator as well as other meso-predators. We collected scats (n = 283) of canid species namely, Wolves, and other predators over a period of 3 years (2014-17). GPS co-ordinates were noted. Scats were analysed with standard scatological techniques to determine diet choices in wolves. Scat samples were washed and examined under the microscope for medullary patterns to identify different prey species based on standard methods (ref). Relative frequencies of occurrence [26] of species were obtained. Biomass consumed was calculated using Consumed mean prey mass (kg) per wolf to excrete one collectable scat as a function of mean prey body mass (x kg) provided per feeding experiment by [27] correction factor 1 (CF1), *y* = 1.798(1−exp(−0.008*x*)) as well as the conventional correction factor, *y* = 0.439 + 0.008*x* [28]. We compared data on diet and perceived depredation by large carnivores as well as other carnivores using χ^2^ test. We also performed correlations to determine relationship between perceived depredation and livestock population.

### DNA verification of select scats

To confirm wolf scats and observer error, DNA extraction from only suspected scat samples (n=118) were done by using commercially available QIAamp stool DNA kit (QIAGEN, Germany) in a dedicated room to avoid contamination with some minor modification. We did not sequence fox scats as fox scats are easily distinguishable due to their small size and quantity. We targeted 148 bp region of the mitochondrial Cytochrome b gene for identifying species using carnivore-specific primers (Ferrell et al, 2000) and all the scat samples were PCR amplified. All PCR reactions were performed on Applied Biosystems thermal cycler (ABI, 2720) in a reaction volume of 10 μl containing 1X PCR buffer, 2.5 mM of MgCl_2_, 200 μM of each d-NTP, 1.25 μg BSA, 4 pM of each primer and 0.25 U of Taq DNA polymerase (Genie) and 1 ul of genomic DNA. The PCR cycling conditions were as follows: initial denaturation at 95°C for 3 mins, followed by 32 cycle of denaturation at 94°C for 30 sec., primer annealing at 55°C for 50 sec. min, primer extension at 72°C for 40 sec. with a final extension at 72°C for 10 min.

Amplified PCR products were cleaned up using Exo-SAP treatment to remove residual oligonucleotides and dNTPs prior to DNA sequencing. Forward and reverse primers of the Cytochrome b gene were used for setting up the cycle sequencing reaction. Unbound ddNTPs were removed by using alcoholic precipitation method and subjected for sequencing to ABI 3130 Genetic Analyzer (Applied Biosystems). Sequence Qualities were determined using Sequence Analysis v 5.2 software (Applied Biosystems) and validated by Sequencer v 4.7 software (www.genecode.com). All good quality sequences were compared with NCBI/GenBank (http://www.ncbi.nlm.nih.gov/) database using BLAST tool and species were confirmed with most homologous sequences (100%) available from NCBI database. Multiple sequence alignment (MSA) was performed using CLUSTAL was implemented in BioEdit v 7.0.9.0 software (Hall 1999). Tracking wolf packs

### Human - Wolf conflict Modelling

We used a presence only modelling Maximum Entropy (Maxent) algorithm, which finds the probability distribution of maximum entropy, that is the most spread out or closest to uniform with limited information about the target species [29–31]. Maxent has a potential to map the spatial distribution of species with fewer locations and has performed well as compared to other available presence only models [32–35].

Wolf presence records were obtained from data of three wolf packs that were GPS collared from a period of 2015-2017. Regular fixes were obtained and presence points were thus used from 2,602 locational fixes that were recorded. A total of 9 environmental layers were used to model wolf presence hotspots. We generated layers from Digital Elevation Model (30 m), Forest cover map (FSI, 2014), Topographic Heterogeneity Map, Ruggedness index, slope. Livestock animal kernel density map were derived from village surveys. Similarly kernel density maps were developed for livestock depredation claims by respondents for wolf and snow leopards. Following layers were then used to create various species distribution models.

i. DEM (30 m) from SRTM data
ii. Topographic Heterogeneity generated from DEM
iii. Forest Cover generated from Forest Survey of India, 2014
iv. Livestock Density Layer – generated from village surveys
v. Ruggedness Layer – Terrain ruggedness index (TRI) is a measurement developed by Riley, et al. (1999)
vi. Slope – generated from DEM
vii. Aspect – generated from DEM
viii. Snow leopard depredation – generated from village survey
ix. Wolf depredation- generated from village survey
x. Wolf scat locations – from field data collection
xi. Wolf habitat use – kernel density layer from GPS telemetry

### Model validation and selection

The predictive performance of the model was estimated using area under the curve (AUC) derived from relative operating characteristic (ROC) plots. To select the best model parameters we compared different models with a combination of the “feature class” and “regularization multiplier”. MaxEnt provides different types of restrictions (“feature class”) in the modelling stage such as lineal (L), quadratic (Q), product (P), threshold (T), and hinge (H). We used all the possible combinations of these features. The used regularization (beta) multiplier values were based on [36] 0.25, 0.5, 1, 2, 3, 4 and 5. Models with lower multipliers tend to over fit data however, higher multipliers may lead loss in capturing species relations to various environmental components [37]. Combining features classes and regularization multipliers, we assessed a total of 42 models for each case study, plus the default auto-feature with 5000 back ground points.

Evaluating whether competing models are significantly different from one another was done in ENMTools (Warren et al. 2010). We compared all the generated models by utilizing the corrected Akaike information criterion (AICc) available in the software ENMTOOLS version 1.4.4 (Warren et al. 2011). The process of choosing the best model using AICc has some limitations that can have an impact on the final results. Results of simple model selection implemented by ENMTOOLS should be carefully interpreted because uncertainties of each parameter values may not be fully incorporated for calculation of AICc. AICc can excessively penalize over parametrization, which can lead to choosing the wrong model (Hastie et al. 2009). Despite this fact, simulation have shown that AICc perform fairly well (Warren & Seifert 2011, Warren et al.2014). We selected the best model and ran 30 replicates with cross validation to test for model consistency and used the averaged best model to finally generate a conflict map in Arc Gis 10.6. The final cumulative output was used to depict conflict values from 0-1.

## Results

### Wolf diet

A total of 118 suspected wolf scats were collected in the study area during the period between 2014-17 (Fig 3). DNA tests confirmed 101 scats that belonged to *Canis lupus* with 99% match in the genbank BLAST database. We found 105 wolf scat samples that were sequenced successfully on amplification. The error rates that were achieved without DNA methods was 85.75%, which means 12.25% of the scats were mis-identified. Out of a total of 118 scats 11% of the scats did not produce any results. Wolf diet constituted mostly of domestic prey (79.02 %) while wild prey constituted to 17.80% of wolf diet over the three years. Remaining prey items which amounted to 3.16% were unidentifiable or made up vegetative matter. In terms of biomass consumed Domestic Cattle contributed the most while rodent species and birds contributed the least (Fig 3).

### Perceived Human-Wolf conflict

Village surveys recorded only 4% of the respondents confirmed wolf presence and perceived them as a possible threat to various livestock (Fig 2 & Fig 4). Over, 98% of the respondents claimed that wolves were not safe for livestock and were averse to its presence. Similarly, 97% of the respondents claimed snow leopards also were a threat to livestock. Claims by 97% of the respondents were that feral dogs as well posed a threat to livestock apart from wolves and snow leopards. We found that villages which reported higher livestock numbers also reported higher livestock depredation (P = 0.47, r = 0.22). Mean livestock depredation cases reported was 5.25 ± 4.71 (S.D), 7.70 ± 9.54 and 8.83 ± 6.05 for wolf, snow leopard and all predators respectively. Claims of wolf depredation showed higher correlation with livestock population in villages (P = 0.67, r= 0.47) than snow leopard depredation (P = 0.56, r = 0.35). We found that perceived level of depredation were significantly different from the actual diet consumed by wolves (χ^2^ = 99.64, p-value <0.0001).

**Fig 2.**
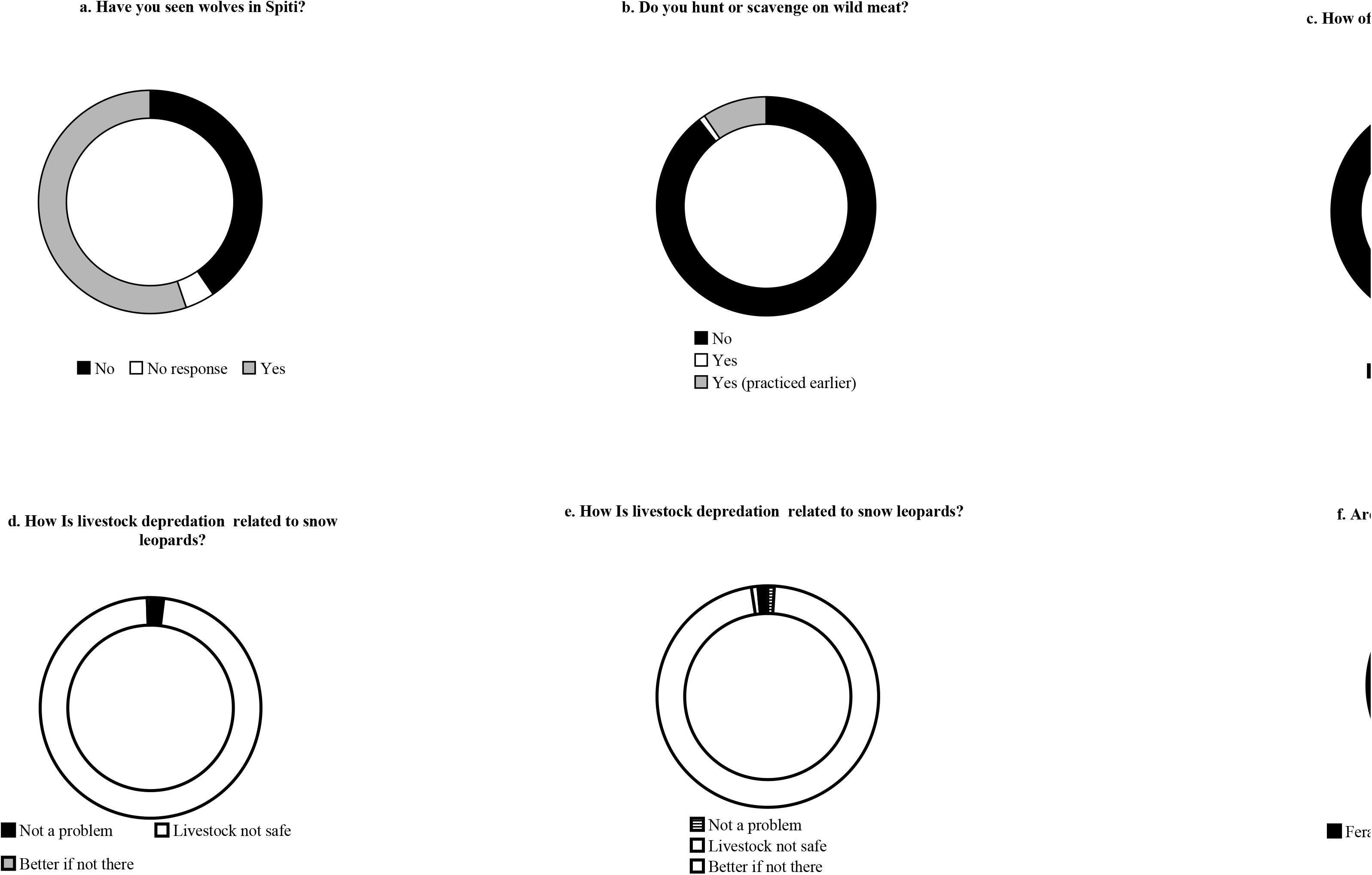
Response with respect to wolf conflict and attitudes of villages in Spiti Region

**Fig 3.**
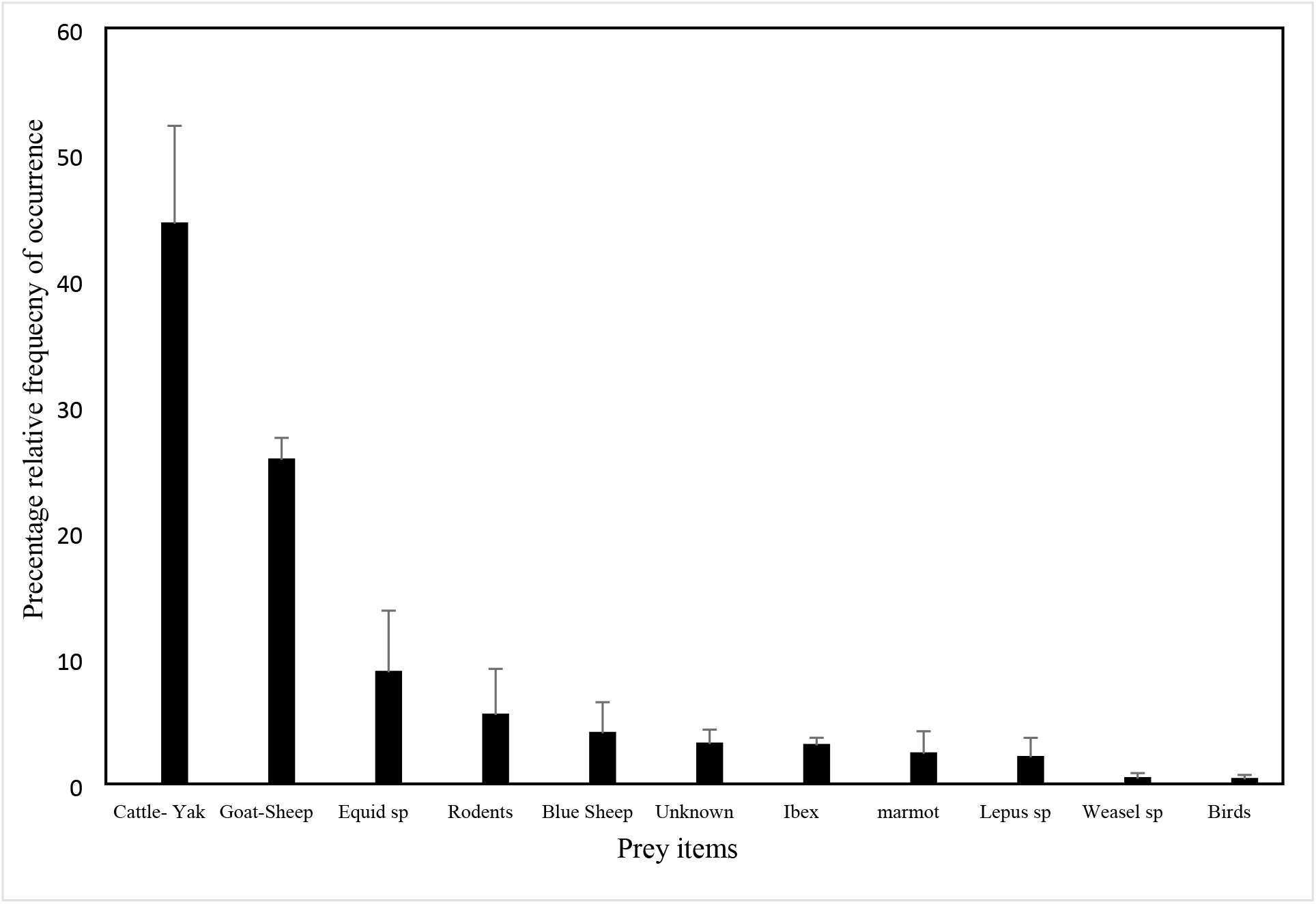
Percentage relative frequency of occurrence of prey items in wolf scats from Spiti region

**Fig 4.**
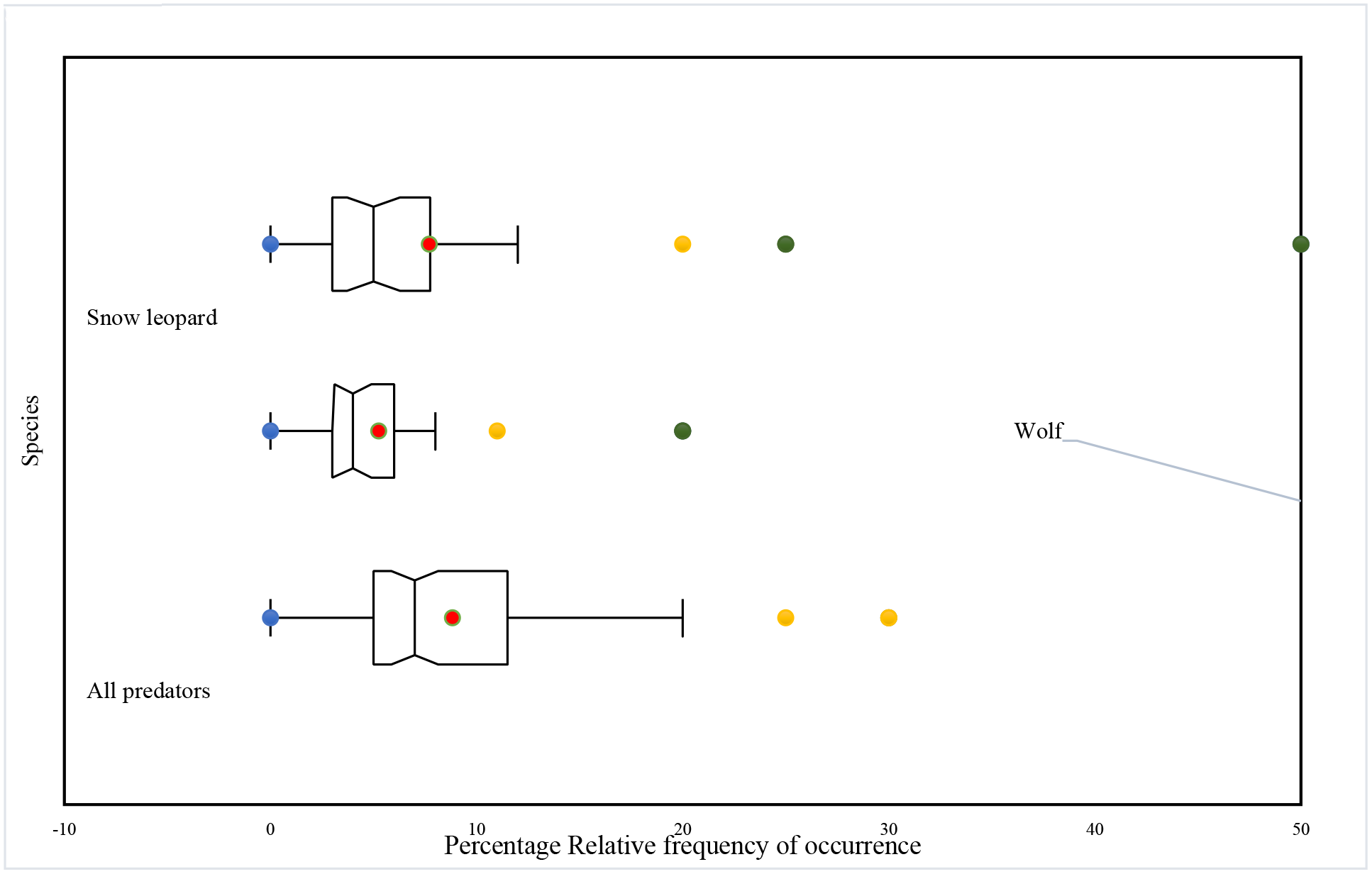
Box Plot of cases of livestock depredation claimed consumed across categories of predators

### Wolf conflict hotspots

Across 42 models we found Auto Model with all features was the best model (beta multiplier = 4, AICc = 2791.86, Fig 5-7). The averaged model output generated a regularized training gain was 1.81, training AUC was 0.94. Test AUC was 0.92 ± 0.04. The average algorithm convergence reported was 227.33 iterations with and average if 5057.03 background points. Wolf depredation (43.65 ± 4.1 %), Animal location (18.07 ± 2.75 %) and Village density (17.54 ± 1.5%) constituted as the top three variables that contributed to AUC. The environmental variable with highest gain when used in isolation was animal location, which had the most useful information by itself. Similarly, decrease in the gain was most when animal location was omitted, therefore generated the most information that was present in the other variables. Marginal response curves depicted the model to have positive responses to animal location, LULC, village population, village density and wolf depredation. Marginal curves were negative to slope, ruggedness and terrain heterogeneity. Curves showed peak like responses for DEM, NDVI and Livestock density. We found area denoted as very high conflict zone was 8 km^2^, high conflict zone was 14 km^2^, medium conflict zone was 61 km^2^, low conflict zone was 91 km^2^ and negligible or no data zone below threshold was 5573 km^2^.

**Fig 5.**
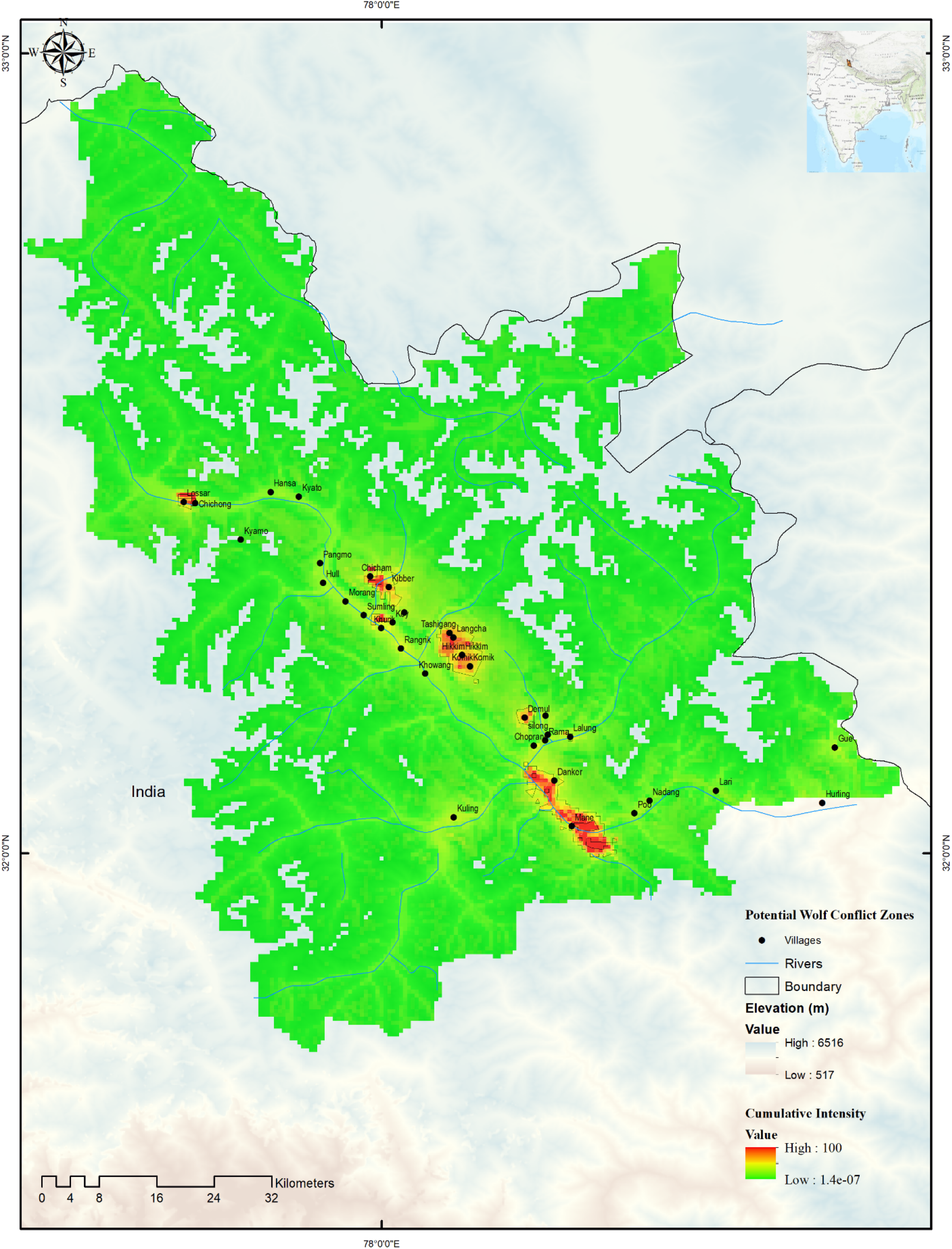
Wolf conflict hotspots modelled. Best model was chosen and hotspots were mapped.

**Fig. 6.**
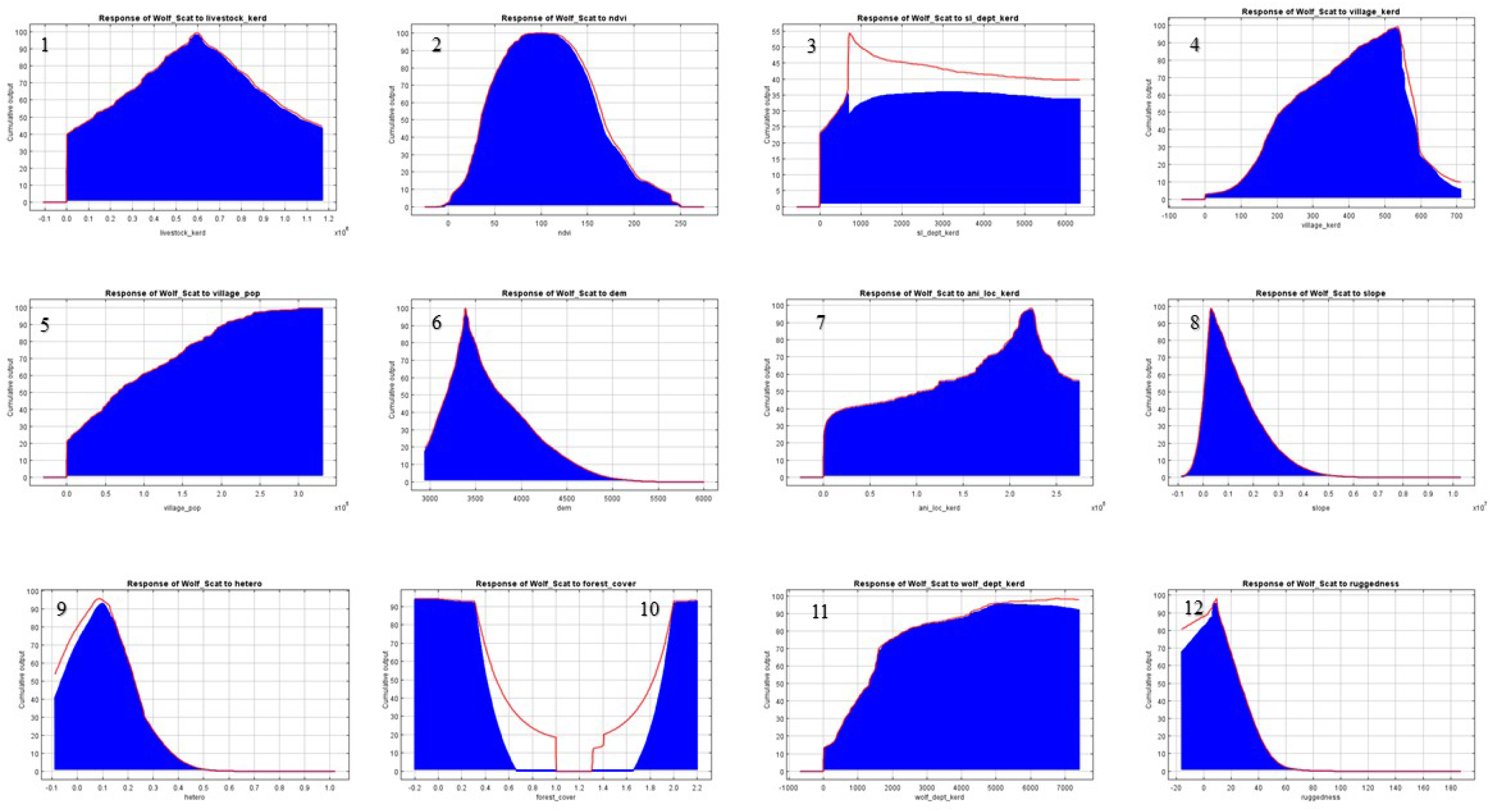
Marginal response curves of variable layers used (n=13). Marginal response curves of LULC not shown as response was did not show any pattern.

**Fig. 7.**
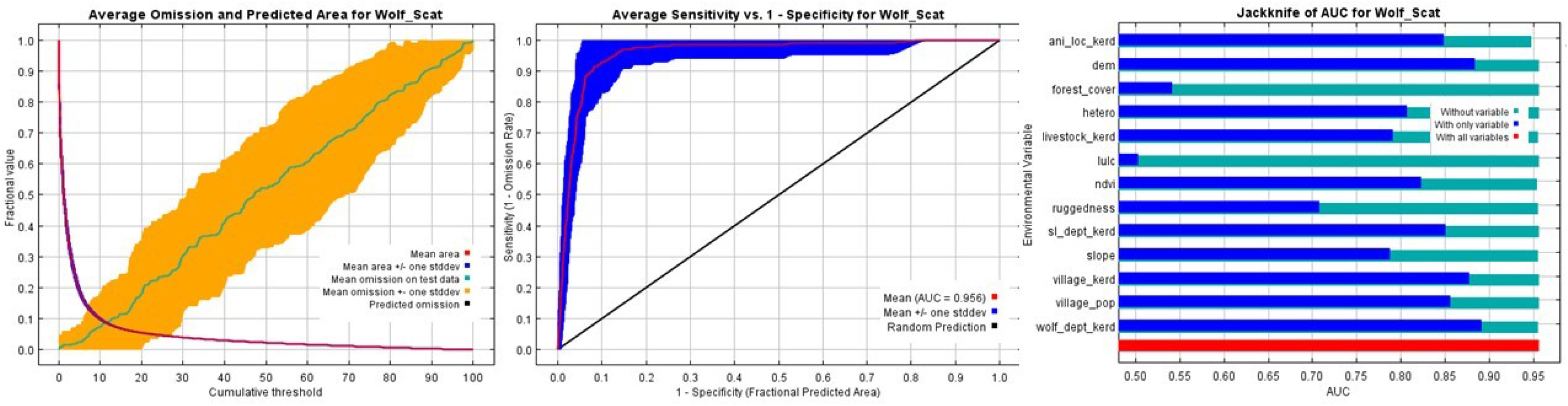
a) Average omission and predicted are of wolf conflict. b) Sensitivity and specificity along with c) jackknife of AUC shown for each variable.

## Discussion

Cultures that raise livestock have strongly disliked wolves [38]. We found that wolves of Spiti largely predated on large domestic prey. Similar studies from the adjoining landscape of the regions reflect comparable patterns [10,16]. A review of dietary habits of grey wolves [13] showed wolves from India preferred mostly medium sized wild ungulates and domestic prey. The results of the review [13] present mostly dietary habits of peninsular wolves of India, the current study thus adds to the existing information. Similarly, Zhang et al. (2009) found that in Dalai Lake Natural Reserve, Mongolia; livestock formed much of the wolf’s diet probably due to overabundance of livestock. These patterns of prey consumption are more in common across central Asia, where Mongolian wolves co-occur in likewise agro-pastorally occupied landscapes (Lyngdoh et al 2019).

Questionnaire surveys and interviews revealed there is a perceived conflict with agro-pastoralists and wolves, which were reported to be one of the main causes of livestock mortality. The northern part of the study area (Kibber WLS and adjacent areas) reported more conflict in terms of wolf predation while the south western part (Near Pin Valley NP area) reported relatively greater conflict arising due to snow leopards. Retaliatory killing of wolves are reported to occur despite unclear perceptions that the snow leopard in responsible for most kills of livestock [39], but this practice has slowly been discontinued largely due to conservation awareness, enforcement as well as religio-social sentiments of the predominant Buddhist communities of the region [11,24]. In comparison, attitudes due to perceived losses due to wolf is similar to examples from North America, Finland [14,38,40].

Our results showed that perceived levels of livestock depredation did not emulate actual levels of depredation in concurrence with earlier studies [12,14]. We found disparities in terms of type of prey and quantity of livestock consumed as claimed by respondents against scatological analysis. Similar findings have been report from the Northern Rocky Mountains where wolf depredation is lower than expected given its exposure to domestic livestock. Wolves accounted for less than 0.04% of the total losses or 0.01% of all predator caused mortalities [38].

The simplicity and ease of use of Maxent has prompted many researchers to use the software [33]. We avoided default settings and approached the modelling process to arrive at an optimal regularization parameter, which may differ if push button approaches are used. The three variables that were important permutation terms were animal location, village density and slope. Scat locations, which correlate, well with animal location were thus a very good indication of conflict hotspots that could serve as reliable data in cases where animal locational data is not available. In terms of percent area of conflict, scale appeared less significant. However, it is important to note that the scat presences correlated well with village population as well. Wolves operated at peaks of 3300 mts with gentle slopes, which was expected. This may be also due to competition with sympatric species such as snow leopard which prefer more rugged and cliff like terrain [41].

Therefore, conflict hotspots reflected were in areas that had low topographic heterogeneity and, decreasing ruggedness. Wolves preferred areas that were undulating and gentler in their slopes. Whether this is driven by competing species or niche exclusion in terms or space and prey is to be examined. Wolves selected area with optimal livestock densities and villages that maintained their rural characteristics. This means it was unlikely to find wolf depredation with increasing urbanization. However, it does not mean that decreasing livestock would reduce chances of wolf presence. Studies have shown that wolves may choose areas wherever wild prey is available in good numbers over areas that were heavily grazed by livestock as well. Predominantly selected areas of wolf depredation were agriculturally inclined village with good livestock and accessible terrain or topography. Wolves also operated at an optimal ndvi range, which could surrogate for availability in terms of wild and domestic prey. From field observations, it is clear that in this landscape of limited and short flowering periods, herds may be concentrated in productive pastures, which are very often occupied by domestic livestock and may lead to prey switching amongst wolves due to sheer likelihood of greater contact, depressed wild prey availability and habit over wolf generations in time.

It is now evident that top predators have capability to drive trophic cascades across large landscapes [42–44]. The wolf is a top predator in this region, its conservation is linked with securing not only the landscape but the people and the region as a whole. In Spiti, a stronghold for Himalayan wolves, cultural ethos and wildlife have been are primary reasons for tolerance to large predators. However, with changing land-use patterns and economic thrust the threshold-tolerance levels for many wild flora and fauna may be altered. Our study therefore provides great understanding of species ecology for furthering conservation actions given the harsh context of the high elevation Himalayan rangelands. The applicability of such methods can be improved through repeated efforts that can aid in policy framework for many Himalayan species. This approach may also aid in informing the management towards timely interventions in order to conserving such an elusive, large ranging and ecologically unique Himalayan wolf.

## Acknowledgements

We thank and appreciate the encouragement given by the Director, VB Mathur and Dean, GS Rawat of WII with this study. We thank specially the Himachal Forest Department, Spiti Wildlife Division, Rajesh Sharma, Rajeev Sharma, Devender Chauhan and Staff for their support. We thank Parag Nigam and S. Krishna for their immense help in immobilization of animals. We thank the help in field by various volunteers and personnel. We thank Gompo, Namtak, Yarphel, Namgial, Dorje, Tenzing, Keysang, Tashi, Rajender and Nishi, Chering, Thukten and their Chammos for their field support. The residents of Mane, Kibber, Langcha, Suluk, Chichong and Kholaksar. We thank the staff at Chandertal. We would like to express our immense gratitude to the people of Spiti for their care and concern during our field times for this study. *Julley!*

